# Indistinguishable network dynamics can emerge from unalike plasticity rules

**DOI:** 10.1101/2023.11.01.565168

**Authors:** Poornima Ramesh, Basile Confavreux, Pedro J. Gonçalves, Tim P. Vogels, Jakob H. Macke

## Abstract

Synaptic plasticity is thought to be critical for building and maintaining brain circuits. Models of plasticity, or plasticity rules, are typically designed by hand, and evaluated based on their ability to elicit similar neuron or circuit properties to ground truth. While this approach has provided crucial insights into plasticity mechanisms, it is limited in its scope by human intuition and cannot identify *all* plasticity mechanisms that are consistent with the empirical data of interest. In other words, focusing on individual hand-crafted rules ignores the potential degeneracy of plasticity mechanisms that explain the same empirical data, and may thus lead to inaccurate experimental predictions. Here, we use an unsupervised, adversarial approach to infer plasticity rules directly from neural activity recordings. We show that even in a simple, idealised network model, many mechanistically different plasticity rules are equally compatible with empirical data. Our results suggest the need for a shift in the study of plasticity rules, considering as many degenerate plasticity mechanisms consistent with data as possible, before formulating experimental predictions.

## 1 Introduction

Synaptic plasticity is the ability of synapses to change their efficacy based on their pre- and postsynaptic environments and it is thought to be critical for the brain’s ability to learn from and remember past experiences [1, 2]. Experimental efforts to understand the “plasticitome”, i.e., the combined action of many synapses [3, 4, 5, 6] are still hampered by the inability to record from multiple synapses simultaneously, *in vivo*, during learning. Theoretical models fill this void, allowing us to link analytical arguments and the available empirical data to propose putative mechanistic plasticity rules to serve a given network function [7, 8, 9, 10, 11].

Critically, such rules rely on human intuition and hand-tuning [12, 13], but they rarely test if more than a *single* plasticity rule could produce a desired network effect (Fig. 1A). Indeed, any possible degeneracy is difficult to explore in hand-tuned systems. On the other hand, degeneracy is so widely observed in neuroscience [14, 15, 16] — and biology more generally — that it would be erroneous to assume singular solutions for a given plastic system.

**Figure 1:**
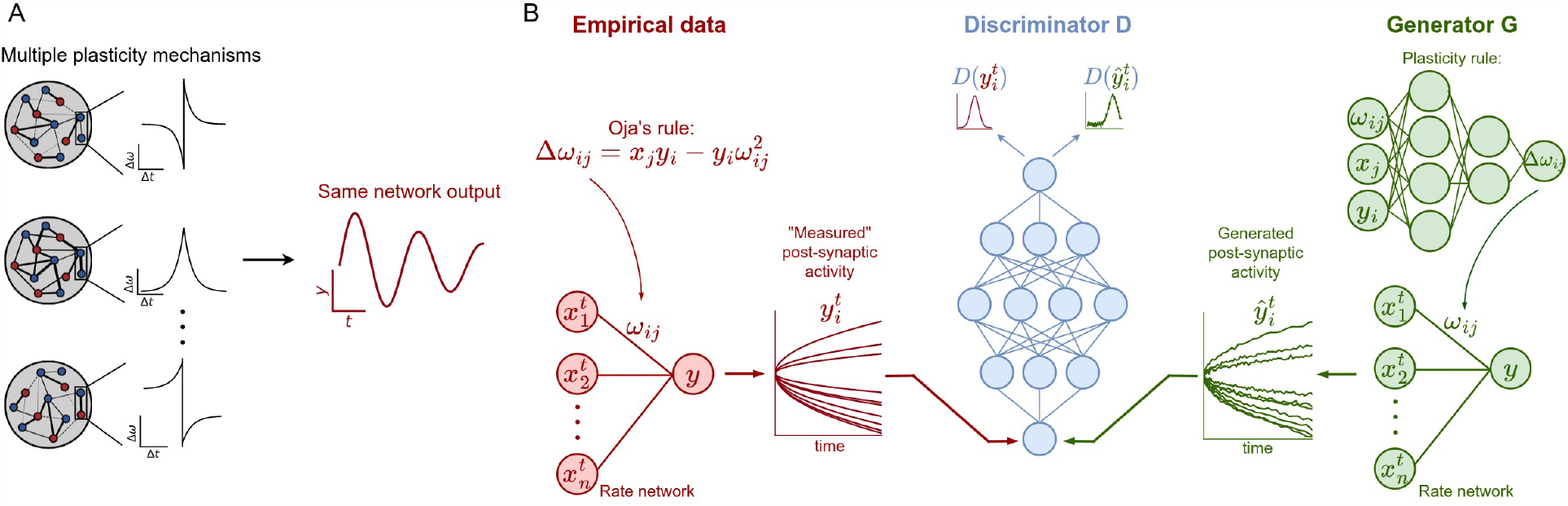
(A) We hypothesise that multiple plasticity mechanisms could lead to the same neural network activities, i.e., there is degeneracy of plasticity mechanisms. (B) Adversarial learning of plasticity rules: empirical data are simulations of the postsynaptic activity of a rate network with plastic synapses evolving according to Oja’s rule. The generator G is a rate network with synapses evolving according to a tunable MLP rule. The discriminator D is a flexible network trained to distinguish empirical data from the generator output. In our framework, the generator and discriminator are trained so that at convergence, the learned MLP rule makes the generator produce postsynaptic neural activity traces indistinguishable from the empirical data.

A promising approach to unveil potential degeneracy in plasticity is to use automated methods to either propose a range of potential plasticity rules for further hand-crafted modelling, or to systematically explore the space of plausible plasticity rules underlying empirical observations or subserving network computations. Such recent attempts to infer plasticity mechanisms with automated methods [17, 18, 19, 20, 21] use flexible parametrizations of plasticity rules, such as Volterra expansions [17, 18, 20] or neural networks [19, 21], to capture the widest possible range of plasticity mechanisms. Handcrafted loss functions (or meta-objectives) are then used to shape the parameters of these rules to satisfy a given biological constraint, e.g., network stability or familiarity detection. Unfortunately, hand-crafted loss functions come with similar issues as hand-crafted rules, in that they can only produce the results within reach of human intuition. For instance, it is difficult to glean a comprehensive loss function from given experimentally recorded neural activity. What’s more, it is near impossible to exclude non-plasticity-related contributions to network activity such as changes in state or input to the network, or measurement variability (Fig. 1A).

Here, we propose to deduce both plasticity rules *and* loss functions directly from empirical data in an adversarial game in which a *generative* Deep Neural Network (DNN) produces plasticity rules that create simulated data good enough to trick a second, *discriminative* DNN into classifying it as empirical. This pair of DNNs, termed Generative Adversarial Networks [22, GANs] attempt to reproduce the statistical properties — rather than achieve a point-to-point match — of a specified dataset.

We hypothesise that this approach allows us to unveil degeneracy in meta-learned plasticity rules while disregarding confounding factors not related to plasticity. Indeed, by construction, GANs enforce a separation between a systematic component due to the plasticity mechanisms and a noisy component due to other confounding factors: since the adversarial game trains the generative DNN to match the distribution of activity traces [22], and the only tunable parameters are associated with the systematic component, i.e., the plasticity mechanisms, the generator is forced to disregard the confounding factors while updating the plasticity rule parameters.

In a proof-of-principle, we show that GANs find multiple, mechanistically different plasticity rules that can produce activity consistent with synthetic empirical dynamics. More specifically, our adversarial approach identifies degenerate plasticity rules on synthetic data simulated with a known plasticity rule, i.e., Oja’s rule [23]. Using synthetic data from a known rule allows us to compare the rules learned with our framework to the ground-truth rule without any unknown additional factors. All the plasticity rules learned by our approach plausibly and robustly generate activity traces that are statistically indistinguishable from the training data, even though their synaptic weight dynamics differ substantially from those produced with Oja’s rule. Our findings point towards a necessary shift from looking for the correct synaptic plasticity rule for a given network function, to inferring entire families of rules with similar network-level function.

## 2 Results

To investigate if multiple plasticity rules could achieve similar neural dynamics, we used an adversarial approach to deduce both the plasticity rules and the loss function from (simulated) *empirical* neural activity. The GAN approach implicitly selects the features of the empirical data [24, 25, 26] relevant to the determine the plasticity mechanisms at play, thus requiring no additional pre-specified constraints on the data (Fig. 1B, but see discussion). We show that our approach reveals many mechanistically different rules consistent with the empirical data, indicating that the plasticity mechanisms are degenerate, even in the idealised setting of our rate networks. Moreover, the recovered rules exhibit similar generalization properties to the ground-truth rule, such as scale-invariance and robustness to measurement noise.

### 2.1 Model set-up and empirical data collection

We considered an idealized setting with a two-layer linear feedforward network with plastic weights (Fig. 1B). We then produced an ensemble of weight- and activity traces using a known plasticity rule, i.e., Oja’s rule [23] (Fig. 2, top row; Sec. Empirical data and Oja’s rule), in which the weights converge to the first principal vector of the input data (Fig. 2C) while the postsynaptic activity assumes the value of the first principal component (Fig. 2B, black traces, Methods). Subsequently, we used this empirical data to train and evaluate our GAN-based solutions. We flexibly parameterized plasticity rules with a multi-layer perceptron (MLP, 3-layers, Methods, Parametrized learning rules). This MLP approximates local plasticity rules, i.e., it updates each synapse in the linear network at timepoint *t* based on the pre-synaptic activity 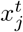, postsynaptic activity 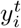 and current synaptic weight of the given synapse 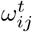. As a control, we confirmed that the plasticity parameterization with the MLP is flexible enough to approximate Oja’s rule, and that the GAN approach is capable of rediscovering Oja’s rule using a constrained search space (Supp. Fig. 5).

**Figure 2:**
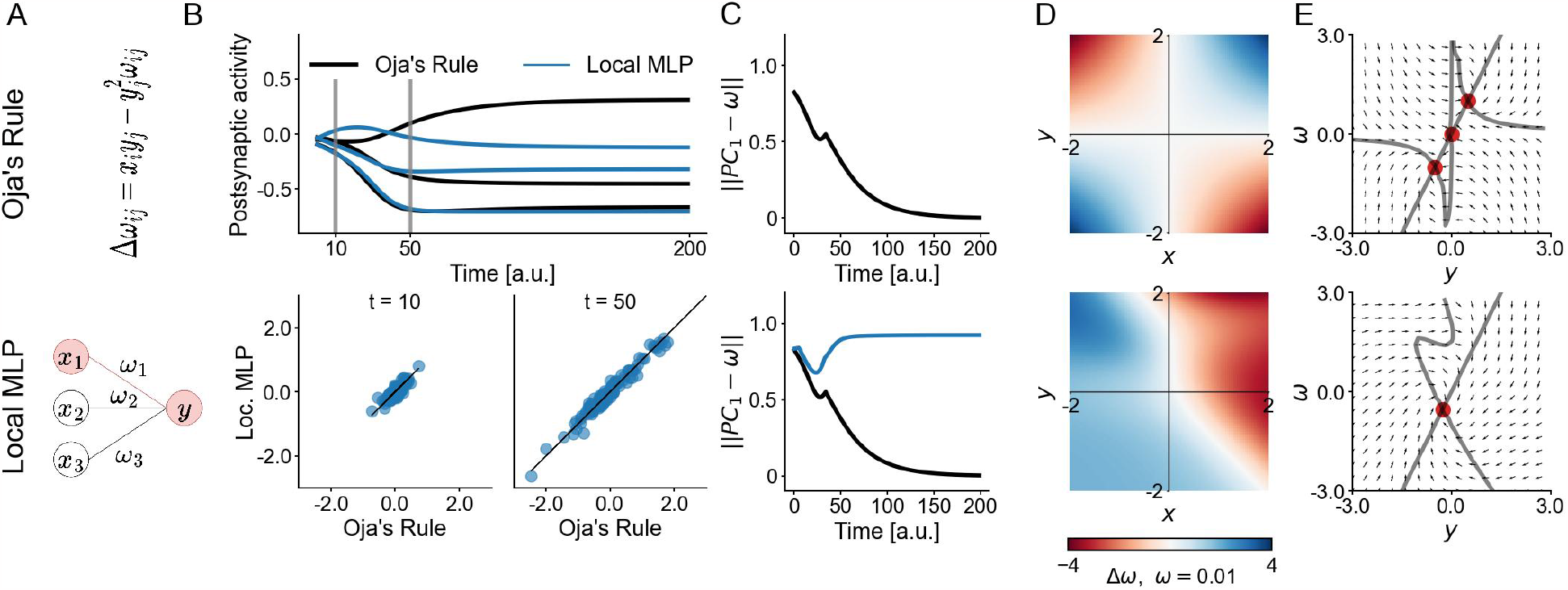
Disparate plasticity rules with same postsynaptic activity. (A) Investigated plasticity rule. (B) Postsynaptic activity traces of a rate network simulated with Oja’s rule (black) and local MLP rule (blue) for different initial synaptic weights (top). Learned-rule activities versus the original Oja’s rule activities at different time points and for different initial synaptic weights (bottom). (C) Weight trajectories, as measured by ||*PC*_1_ − *ω* || . Oja’s rule (top), local MLP (bottom). (D) Synaptic weight updates Δ*ω* for a range of presynaptic activities *x* and postsynaptic activities *y* and *ω* = 0.01. Oja’s rule (top), local MLP (bottom). (E) Vector field of *ω* versus postsynaptic activity *y* with presynaptic activity fixed at *x* = 0.5 (nullclines in black, fixed points in red). Oja’s rule (top), local MLP (bottom).

### 2.2 Disparate plasticity rules with same postsynaptic activity

We began our investigation by adversarially training the MLP-rule on a large collection of (synthetic) empirical data (Fig. 2, bottom row). The thus-trained MLP-rule elicited activity traces that qualitatively captured the salient features of the empirical postsynaptic activity, e.g., the transient increase in postsynaptic activity at earlier timepoints, and the convergence to a stable value at later time points (Supp. Fig. 6). Importantly, the MLP rule reproduced the statistics of held-out empirical data (Fig 2B, bottom panels), even when single trials diverged from the empirical activity traces, in line with our expectation that GANs train on the ensemble rather than on the individual trials.

However, the evolution of the synaptic weights dictated by the trained MLP-rule diverged from the trajectories predicted by Oja’s rule (Fig. 2C), as quantified by the Euclidean distance between the synaptic weights and the first principal component of the pre-synaptic activity PC_1_ at each time point [17]. As expected this distance decayed to 0 for Oja’s rule [23] but not for the trained MLP-rule (Fig 2C), indicating that although the network activity was similar, the synaptic weights evolving with the trained MLP rule never converged to PC_1_.

We could also compare the ground-truth and trained MLP rules more systematically by computing the hypothetical synaptic weight update for a range of presynaptic *x*_*j*_ and postsynaptic *y*_*i*_ activities — the observable network variables — at the fixed synaptic weight *ω*_*ij*_ = 0.01. The resulting heatmap of synaptic weight-updates Δ*ω*_*ij*_ = *f* (*x*_*j*_, *y*_*i*_, *ω*_*ij*_ = 0.01) shows a large qualitative difference between Oja’s rule (used to produce the empirical data) and the trained MLP rule, both regarding the magnitude of weight updates and the ranges of pre- and post-synaptic firing rates leading to potentiation (Δ*ω*_*ij*_ *>* 0) and depression (Δ*ω*_*ij*_ *<* 0) (Fig. 2D). For instance, Oja’s rule is symmetric along the diagonal and anti-diagonal, whereas no such symmetry could be observed in the trained MLP rule.

In order to further compare the interaction of synaptic weights and post-synaptic activity of Oja’s and the MLP rule, we computed the vector-field of the postsynaptic activity and the synaptic weight for a fixed pre-synaptic activity of 0.5 (Fig. 2E). Plotting this vector-field allowed us to understand how the coupled dynamical system of weights and network activity evolved, and thus compare different rules. Like for other measures, the weight update diagrams and their subsequent dynamics of the MLP rule showed large differences from Oja’s rule. Interestingly, the trained MLP rule had only one stable fixed point. Collectively, our results suggest that there exists at least one plasticity rule that is mechanistically different from Oja’s rule but produces statistically indistinguishable activity.

### 2.3 Learned rules with same generalisation properties as Oja’s rule

Plasticity is only one of many neural mechanisms underlying network dynamics. Thus, it is plausible that two networks with the same plasticity rule have different recorded dynamics, due to a plethora of other sources of variability such as changing inputs, noisy individual neuron dynamics, and noisy measurements. We wondered whether our GAN approach is able to ignore contributions to neural variability from sources unrelated to plasticity.

In order to address this, we generated data from two networks with Oja’s rule, with different settings compared to the original network: additive noise in the post-synaptic neuron (Fig 3A); and increased number of pre-synaptic neurons (from 3 to 39, Fig 3B, see Methods for details). We trained a local MLP rule with the GAN approach on each of these two different datasets. We then used the two resulting MLP rules to generate post-synaptic activity in the original setting, i.e., rate networks with only 3 pre-synaptic neurons and noiseless post-synaptic activity. Note that the rules were *not* trained on the original setting.

**Figure 3:**
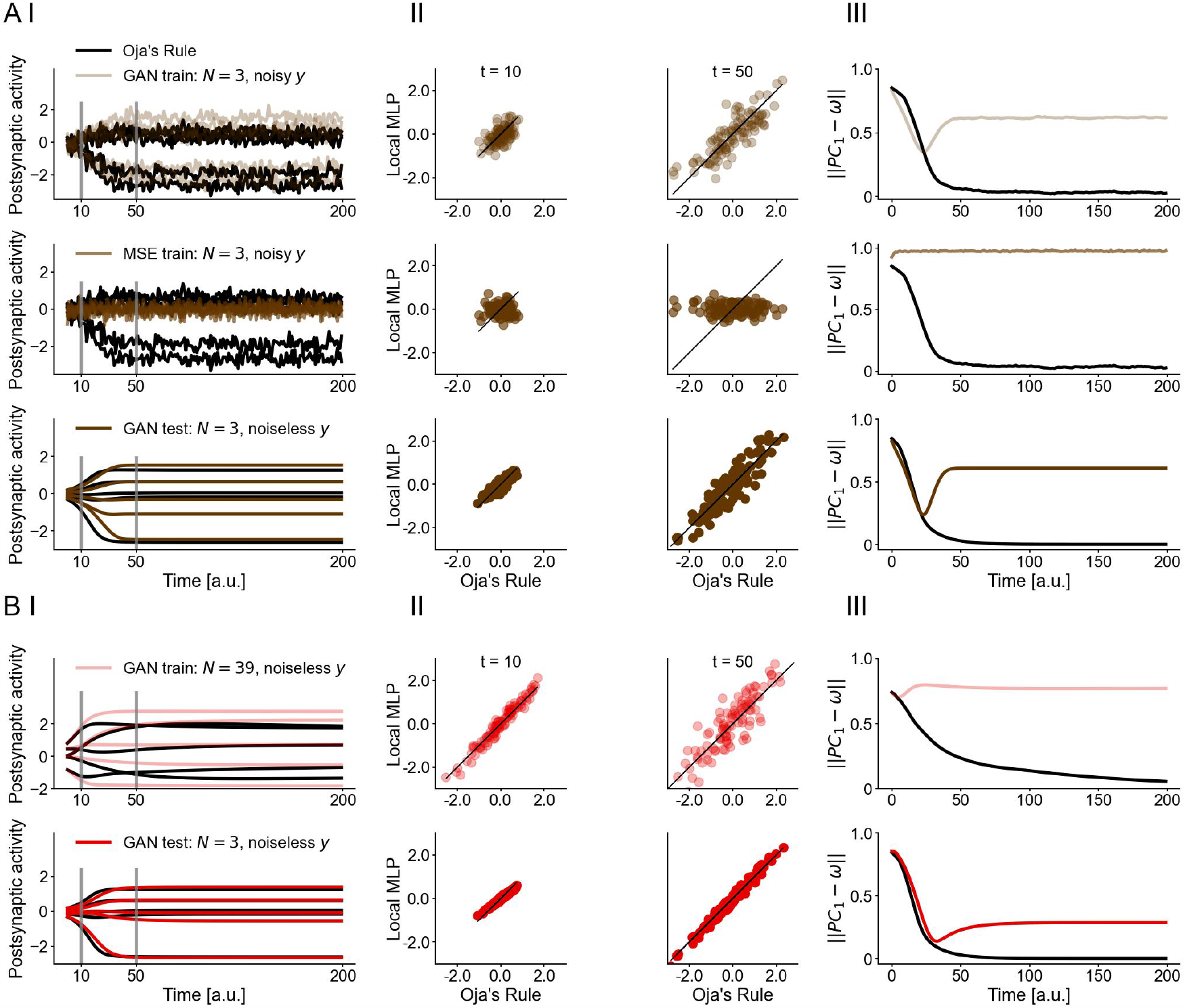
Learned rules have same generalisation properties as Oja’s rule. (A) MLP rule on 3 presynaptic neurons and a noisy postsynaptic neuron: trained with GAN and tested on the same network (top, light brown), trained with a mean-squared error loss and tested on the same network (middle, brown), and trained with GAN and tested on a network with 3 presynaptic neurons and a noiseless postsynaptic neuron (bottom, dark brown). (B) MLP rule on 39 presynaptic neurons and a noiseless postsynaptic neuron: trained with GAN and tested on the same network (top, pink) and on a network with 3 presynaptic neurons and a noiseless postsynaptic neuron (bottom, red). (I) Trajectories of postsynaptic activity for various synaptic weight initialisations generated with GAN-learned MLP rules are qualitatively similar to those from Oja’s rule. (II) Activites from GAN-learned rule at different time points match the statistics of Oja’s rule for both held-out data from the training network and test network. (III) Weight trajectories for learned plasticity rules. Oja’s rule in black.

We found that the two MLP rules, specifically the *same* rules that captured the activity statistics of the training data, accurately reproduced the statistics of the empirical activity traces from the original test dataset, and therefore successfully ignored the perturbations included in both training datasets (Fig. 3 I, II). As before, the evolution of the weights of the two learned rules and their vector-field dynamics differed substantially from Oja’s rule (Fig. 3III and Supp. Fig. 7). In addition, the ability of our GAN-learned MLP rules to generalise over different datasets such as the perturbed datasets used above was not matched by MLP rules trained with supervised loss functions (Fig. 3A, Mean Squared Error on the activity trajectories), highlighting again that selecting the features of the data, and by extension, crafting a loss function to fit plasticity — and avoid overfitting — is crucial and non-trivial.

Collectively, these findings confirm that multiple, mechanistically different plasticity rules can shape networks to elicit similar activity as Oja’s rule. Moreover, our adversarially-trained plasticity rules exhibit remarkable generalization properties between disparate training data and succeed in retaining only the features from the training data that are directly connected to the plasticity mechanisms. In other words, the variety of learned rules we observed stems from a degeneracy in plausible plasticity mechanisms and not from other sources of variability in the data.

### 2.4 Different learned rules due to modeller bias

Next we considered the effect of training different classes of plasticity rules on the same empirical data. So far, we used a parametrization that relied exclusively on local synaptic information, i.e., pre- and post-synaptic activity, and the synaptic weight. We expanded the parametrization so that the weight-update was also dependent on the average pre-synaptic activity and average synaptic weight of all synapses at the time of the update. We called this parametrization “semi global”. In addition, we considered a “global” parametrization in which all neurons’ rates and all synaptic weights were available to every synaptic weight update. Finally, we considered a local MLP biased towards Oja’s rule (see Methods).

As before, the trained rules reproduced the statistics of the empirical data (Fig 4B), while their weight dynamics differed from Oja’s rule (Fig 4C). Notably, they also differed from the weight dynamics of the previously learned rules (Fig 4D,E), showing that different rule parametrizations lead to different learned rules, and thus suggesting the existence of several optimum points in the landscape of learning rules that are all equally consistent with the empirical data. Varying model architectures may nudge the parametrized rule closer to one of these points in the rule landscape, resulting in convergence to a different plasticity rule depending on the model architecture.

**Figure 4:**
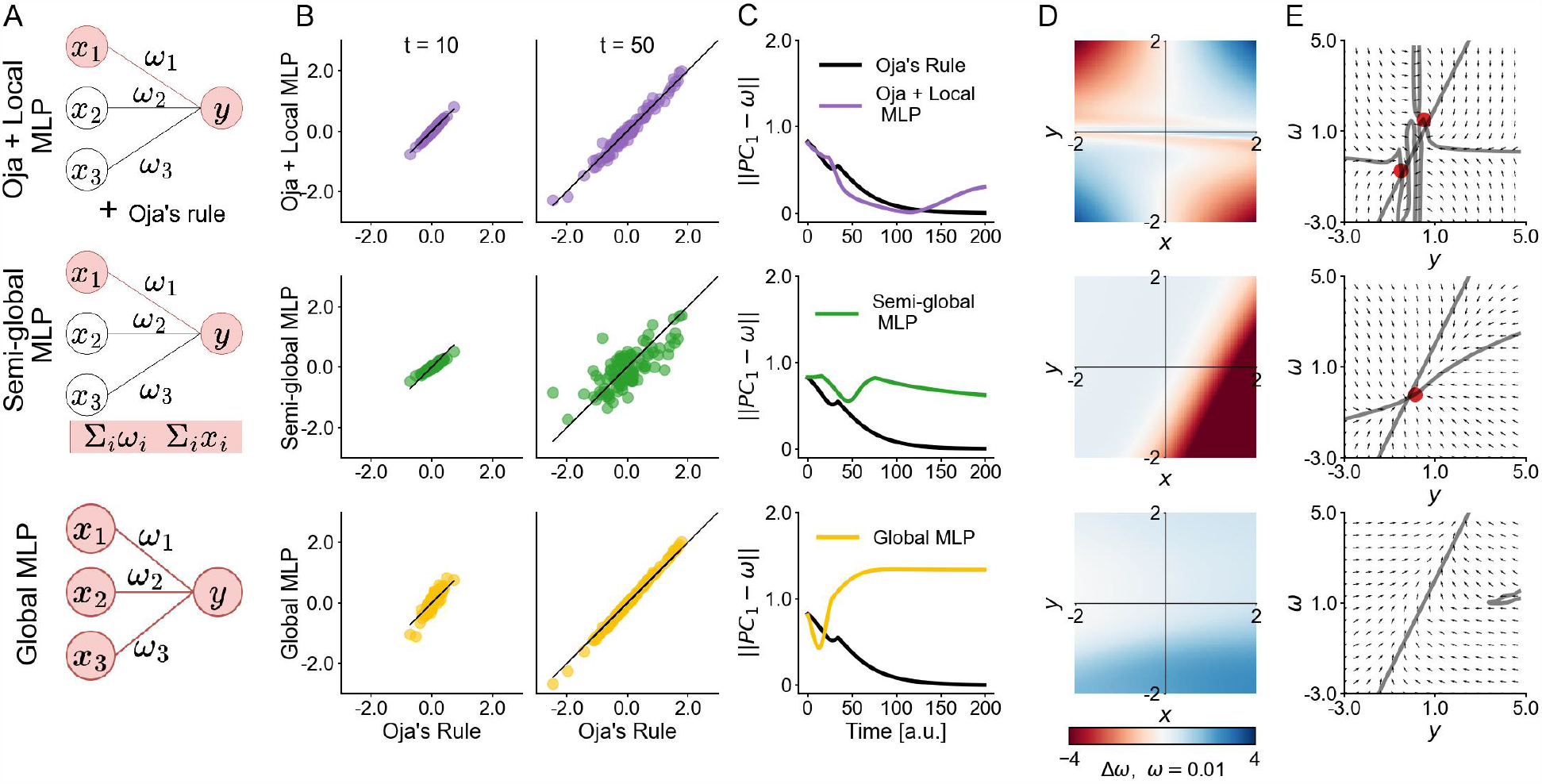
Different learned rules due to modeller bias. (A) Parametrized plasticity rules. Oja+Local MLP (top), semi-global MLP (middle) and global MLP (bottom). (B) Learned-rule activities versus the original Oja’s rule activities at different time points and for different initial synaptic weights. (C) Weight trajectories, as measured by ||*PC*_1_ – *ω* || for Oja + local MLP (top, purple), semi-global MLP (middle, green), global MLP (bottom, yellow). (D) Synaptic weight updates Δ*ω* for a range of presynaptic activities *x* and postsynaptic activities *y* and *ω* = 0.01. (E) Vector field of *ω* versus postsynaptic activity *y* with presynaptic activity fixed at *x* = 0.5.

## 3 Discussion

Theoretical studies about synaptic plasticity typically postulate specific, biologically plausible network functions and hand-design rules to produce them. They often focus on small numbers of canonical rules, with a few terms capturing first-order dependencies on synaptic variables [27, 8, 11]. However, each specific rule provides only one of potentially many consistent explanations for a given dataset, and might overfit to confounding factors inherent to the empirical data. Without exploring the space of possible solutions, it is difficult to assess the validity, completeness, and robustness of a given rule, or to judge its role in memory and learning.

Here, we introduced a GAN framework that allowed us to flexibly explore a larger and less biased space of solutions by algorithmically identifying plasticity rules directly from data. By adversarially training the plasticity rule and the loss function, we found a multitude of rules that captured the same neural dynamics. Crucially, these GAN-learned rules were not variations of the same rule with multiplicative or additive factors, but exhibited a wide range of different weight dynamics and fixed points. Furthermore, these novel rules also differed depending on the rule parametrizations, i.e., modeller bias. Finally, we showed that under subsampling conditions or added observation noise, the inferred rules still captured the statistics of the original unperturbed test dataset. This robustness to different sources of variability in the empirical data indicated that the underlying plasticity mechanisms were truly degenerate.

Degeneracy is present all throughout biology, and more specifically in neuroscience [14], and it would be surprising if synaptic plasticity mechanisms were the exception. Our study shows that degeneracy is present even in idealized systems, and that learned plasticity rules are strongly influenced by modeller bias. Our results suggest that we must account for degeneracy in plasticity mechanisms in order to gain meaningful insights about their role in neural computations. However, the theoretical possibility of degenerate solutions as shown here does not prove their biological implementation, and several more steps are necessary to provide experimentally tractable rules and testable predictions. Since the number of observable neurons in a given experiment is much smaller than the number of synaptic variables in the system, constraining plasticity rules by neural activity measurements alone may not resolve the issue. It will thus be helpful to build more biologically detailed models to make predictions on what experimental features to focus our attention on. Alternatively, some of these issues might be resolved by adopting probabilistic machine learning approaches to obtain uncertainty estimates for plasticity rules inferred from neural activity [28], although this remains a subject for future work.

Our investigations focused solely on Oja’s rule: while this is a canonical rule which has been extensively studied, it is likely too simple to form a realistic representation of plasticity mechanisms in the brain. In principle, it is conceivable that more complex rules [12, 13], operating in larger neural populations implementing more sophisticated computations, could be inferred from data unambiguously. However, it is possible (and we would argue probable) that adding complexity in the system will make degeneracy even more likely.

We chose GANs because of their ability to automatically and implicitly select features from data, thus removing the need to hand-craft loss functions. In addition, since GANs attempt to capture *distributions* of activity traces rather than specific traces, the learned rules are largely agnostic to trial-specific features which are uncontrolled for in the experimental setting and independent of synaptic plasticity, such as e.g., the attentional state of an animal [29], or experimental noise. However, despite the flexibility to learn synaptic plasticity rules that GANs enable, their application can be technically demanding: they are notoriously hard to train, sensitive to initial conditions, and prone to mode collapse [30, 31, 32]. Furthermore, GANs only provide one consistent plasticity rule per training run, such that full exploration of the space of solutions is arduous. How this approach scales to more complex, higher-dimensional systems is a question for future work.

## Conclusion

Degeneracy is an ubiquitous phenomenon in neuroscience, and its implications are crucial for our understanding of neural computation in general and the mechanisms of plasticity in particular. Our attempt to flexibly learn rules directly from data in a simple scenario reveals that different plasticity rules can explain the same data equally well, provided the generative model of plasticity is expressive enough. This suggests we should shift the way we think about plasticity rules: not as a singular mechanism implemented by every synapse, but rather as families of rules with similar network-level function and with potential mechanistic differences across different synapses. And rather than studying one rule at a time, one might attempt to characterize properties across ranges of rules, and potentially derive predictions *shared* across plasticity rules. These predictions are likely to be more robust than predictions idiosyncratic to a specific rule, and also suggest a more topological view of plasticity.

## Acknowledgements

We thank Everton J. Agnes, Friedemann Zenke, Chaintanya Chintaluri, Auguste Schulz, Richard Gao and Julius Vetter for helpful discussions and feedback on the manuscript. This work was funded by the German Research Foundation (DFG; Germany’s Excellence Strategy MLCoE – EXC number 2064/1 PN 390727645), the German Federal Ministry of Education and Research (BMBF; Tübingen AI Center, FKZ: 01IS18039A) and the European Reseach Council (ERC consolidator grant SYNAPSEEK).

## 4 Methods

Below, we introduce the rate network model, Oja’s rule: the rule underlying our empirical (synthetic) data, the different parametrizations of the learning rules and the GAN-based meta-learning framework.

### 4.1 Network model

We consider a linear feedforward rate network with *N* presynaptic neurons with activity *x*_*j*_(*j* = 1 … *N*), *M* postsynaptic neurons with activity *y*_*i*_(*i* = 1 … *M*), and synaptic weights *ω*_*ij*_. The postsynaptic activity at (discretized) time *t* is updated as follows:

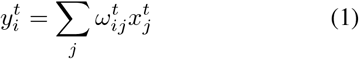

### 4.2. Empirical data and Oja’s rule

In lieu of experimental data, and to test our framework, empirical data consisted of activity traces from the rate network defined above, evolving with Oja’s rule. This rule consists of a Hebbian term, with an added normalization term for stability:

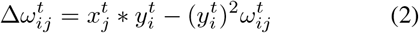

This Hebbian plasticity rule, when used to update the synaptic weights *ω*_*ij*_ of the feedforward rate network in Eqn 1, causes *ω*_*ij*_ to converge to the first principal component PC_1_ of the presynaptic activity 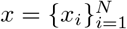 [23]. Concurrently, under Oja’s rule, the postsynaptic activity *y* converges to a projection of *x* onto its first principal component. In order to simulate activity traces *y*_*i*_, we sampled presynaptic activity *x* from a N-dimensional Gaussian distribution, with a given covariance structure (details in Supp. Sec. B.1). Note that we fixed the presynaptic activity to be constant across time, and that the synaptic weight update Δ*ω*_*ij*_ at each time step was computed by averaging over an ensemble of pre- and postsynaptic activities corresponding to *K* = 100 different *x* samples from the multivariate Gaussian distribution. In other words, we have implicit batch learning for the synaptic weights:

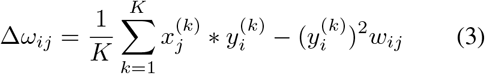

We simulated the plastic network for *T* = 200 time steps. The empirical data consisted of the postsynaptic activity at every time step for each of *K* = 100 different 3-dimensional presynaptic activities and randomly initialised synaptic weights 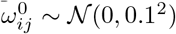.

In Fig 3, we used two other types of data: (1) we introduced noise in the postsynaptic activity: 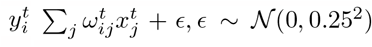; (2) 39 input neurons instead of 3 were simulated, the training was otherwise similar.

### 4.3 Parametrized learning rules

In order to meta-learn the synaptic plasticity rules from the empirical data, we formalized the learning rule with a parametrized function *h*_*θ*_:

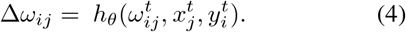

First, for a proof of principle (Supp. Fig. 5), we parametrized Oja’s rule with learnable coefficients *θ*_1_ and *θ*_2_:

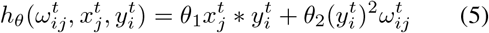

For all other experiments, the plasticity rules were parametrized using multi-layer perceptrons [MLPs, 33], with different inputs from the rate network, depending on the chosen model:

- *Local MLP*: the plasticity rule is parametrized by a 3-layer MLP, i.e., 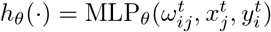 This MLP represents a *local update*, i.e., it transforms each 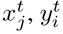 and 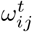 in the same way, independently of the indices *i, j* and *t*.
- *Oja* + *local MLP*: 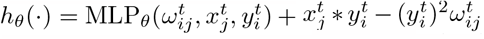 . This learning rule is “biased” since, by construction, it is initialised close to the ground-truth solution and any non-zero outputs of the MLP are perturbations to Oja’s rule.
- *Semi-global MLP* computes the synaptic weight update 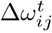 for a single synapse. It takes into account the mean presynaptic activity and the mean across the network synaptic weights at the current time step, in addition to the local presynaptic activity 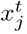, synaptic weight 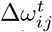 and postsynaptic activity 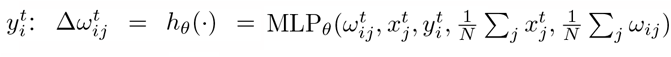.
- *Global MLP* takes into account all pre- and postsynaptic activities and synaptic weights: 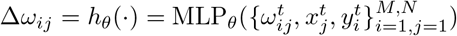.

Note that these parametrized rules are progressively less constrained, and that each MLP could, in principle, be reduced to the one above it.

### 4.4. Meta-learning framework

Our meta-learning framework uses a GAN to learn the parameters *θ* of the plasticity rule *h*_*θ*_, given neural activity. Below, we first introduce the GAN formalism, and then show how one can recast the problem of meta-learning within this formalism.

GANs are a machine learning approach to obtain generative models of data, by training a model to match a target distribution *p*(*d*^*t*^), which we only have access to via samples *d*^*t*^ from the distribution. GANs consist of two deep neural networks: a generator network *g*_*θ*_ that produces data *d* = *g*_*θ*_(*z*), by deterministically transforming latent random variables *z* sampled from a known distribution *p*(*z*). The second network is a discriminator D_*Ψ*_ which aims to classify generated samples *d* as fake (i.e., from the generator), and *d*^*t*^ as real (i.e., from the target distribution, Fig. 1B). After the two networks have been trained with a minimax loss

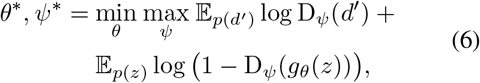

in the limit of infinite data, the generator implicitly represents *p*(*d*^*t*^) at convergence. Thus, convergence yields a generative model of *d*^*t*^. Note that the GAN does not impose any restrictions on the architecture of the generator network. Furthermore, it does not define an explicit loss function for the target data *d*^*t*^ and generated data *d*: instead, the discriminator implicitly represents a distance function between the two data distributions, and this function is also learned end-to-end with the generator. This leads to GANs not attempting to reproduce the data *d*^*t*^ in minute detail, but rather to capture the general statistics of the data. In other words, the GAN matches the distributions *p*(*d*) and *p*(*d*^*t*^), rather than individual samples *d* and *d*^*t*^.

This flexibility in the GAN framework is advantageous for meta-learning plasticity rules from neural activity. We first recast the system composed of the postsynaptic update (Eqn 1) and the plasticity rule (Eqn 4) as a generative model of postsynaptic activity traces for *T* time steps, i.e., 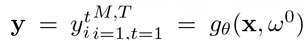, where 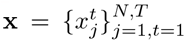and 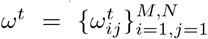 (Fig. 1B). We then define a discriminator D_*Ψ*_ that differentiates between generated activity traces y and empirical activity traces y^*t*^ *∼p*(y^*t*^). Note that y is a random variable, since the rate network is initialised randomly before every forward pass from the generative model. We learn the parameters of the plasticity rule *θ* and the discriminator *Ψ* using the same minimax loss as in Eqn 6:

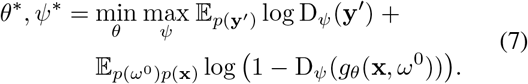

At convergence, the GAN will have learned a plasticity rule *h*_*θ*_ that is consistent with the empirical neural activity **y**^*t*^. The GAN framework thus allows us to flexibly parametrize the plasticity rule. It also allows us to learn the rule from neural activity, without having to specify a loss function on the neural activity. Details on method implementation and numerical experiments are in Supp. Sec. B.

## Footnotes

To lighten the notation, we elide the dependence on time *t* in this equation.

