## Supplementary material for "Indistinguishable network dynamics can emerge from unalike plasticity rules"

### A Supplementary figures

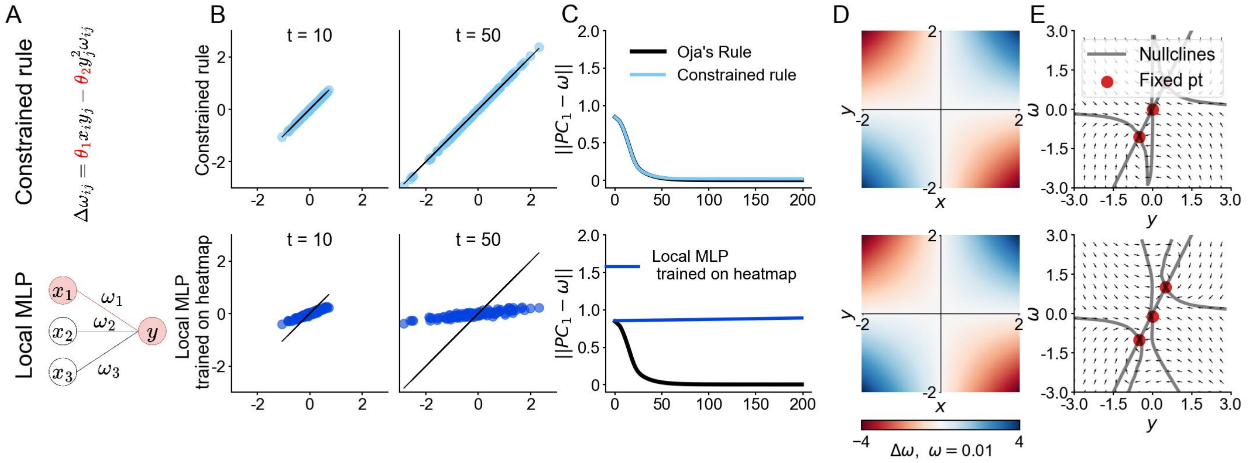


Figure 5: **GAN with constrained rule and MLP rediscover Oja's rule** Oja's rule (black), constrained rule (top, light blue), MLP (bottom, dark blue). (A) Parametrized plasticity rules: Constrined rule (top) and local MLP trained directly on pre-post activity heatmap (bottom). (B) Learned-rule activities versus the original Oja's rule activities at different time points and for different initial synaptic weights (top and bottom). (C) Distance between the synaptic weights $\omega$ and the first principal component of $\mathbf{x}$ across time. (D) Synaptic weight updates $\Delta\omega$ for a range of presynaptic activities $x$ and postsynaptic activities $y$ and $\omega = 0.01$. (E) Vector field of $\omega$ versus postsynaptic activity $y$ with presynaptic activity fixed at $x = 0.5$.

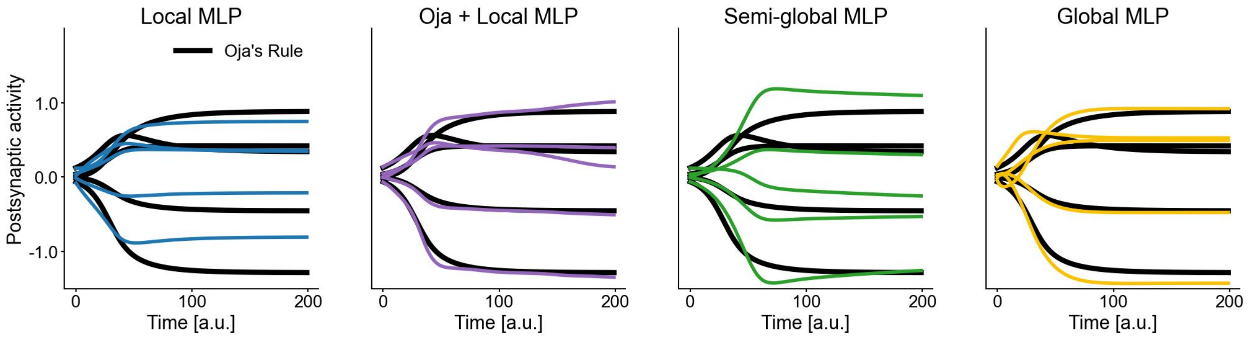


Figure 6: **Activity generated from learned rules captures the salient features of postsynaptic activity traces from Oja's rule.** Postsynaptic activity traces for different initial synaptic weights in the rate network from Oja's rule (black) and learned rules.

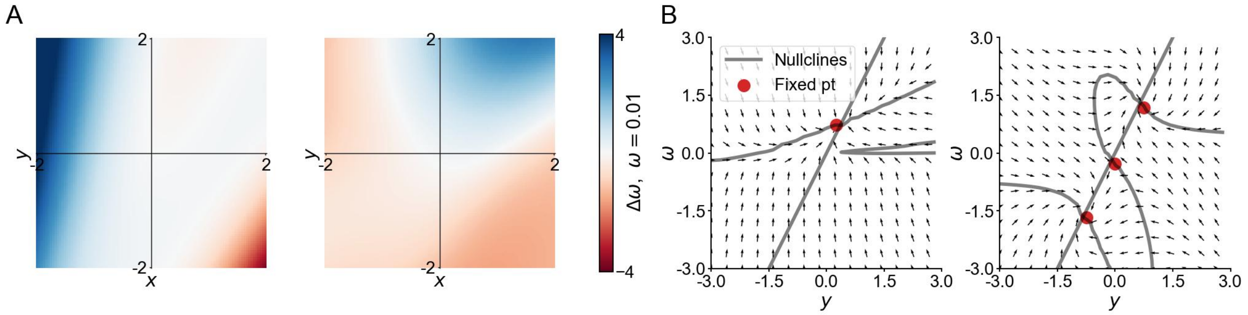


Figure 7: **Dynamics of GAN-based MLP rules trained on perturbed data also differ from Oja's rule.** Local MLP trained on noisy postsynaptic neuron with 3 presynaptic neurons (left); local MLP trained on noiseless postsynaptic neuron with 39 presynaptic neurons (right). (A) Weight updates $\Delta\omega$ for the learned rules are different from that of

Oja's rule. (B) Vector fields of $\omega$ versus postsynaptic activity $y$ for learned rules. Left: 1 stable fixed point; top: 2 stable fixed points and 1 saddle node (middle).

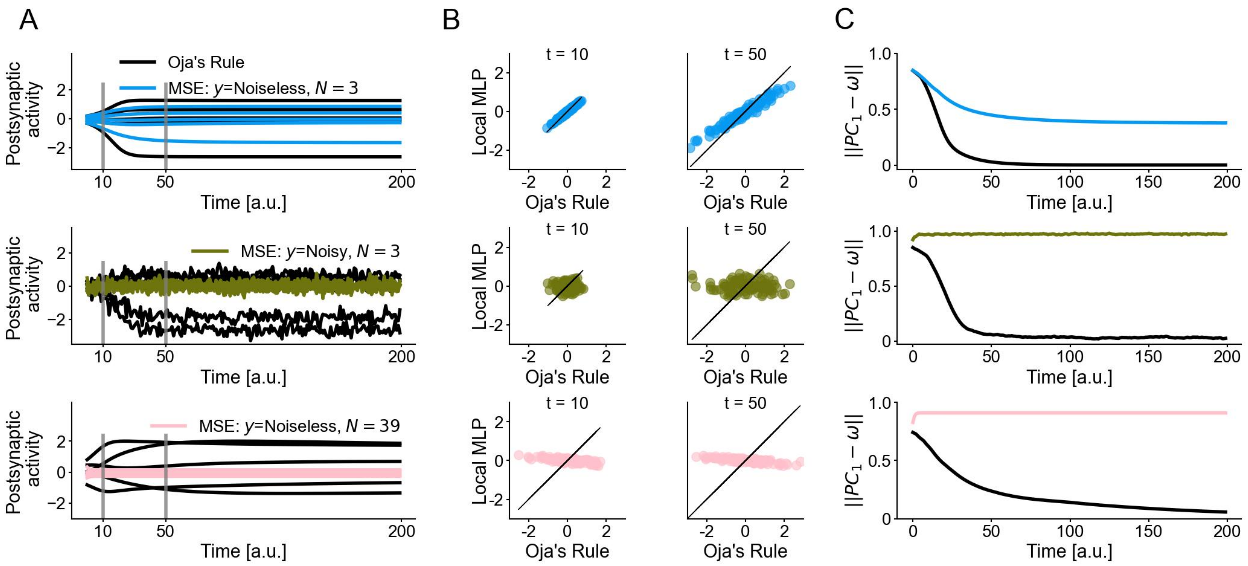


Figure 8: **MLPs trained using a mean-squared loss on the post-synaptic activity do not capture the statistics of groundtruth data.** MLP trained with 3-presynaptic neuron noiseless rate network (top, blue), trained with 3-presynaptic neuron rate network with noise added to post-synaptic activity (middle, brown) and trained with 39-presynaptic neuron noiseless rate network (bottom, pink). (A) Learned rules do not capture the statistics of the postsynaptic activity traces generated with Oja's rule. (B) Weight trajectories from learned rules do not match that of Oja's rule (black). (C) Synaptic weight updates differ from Oja's rule

### B Implementation details

We implemented the network model and GANs in the PyTorch framework [34]. We used Weights and Biases [35] to log different GAN training experiments.

#### B.1 Network model implementation

The network model was feedforward linear, with $N$ presynaptic neurons and $M$ postsynaptic neuron, and thus, $M \times N$ synaptic weights. The network received as input the presynaptic activity $\mathbf{x}$, a $K \times N$-dimensional vector: $K$ samples from a $N$-D Gaussian distribution with a specified covariance matrix $\Sigma$ i.e., $\mathbf{x} \sim \mathcal{N}(0, \Sigma)$. We specified the covariance matrix as follows:

- We constructed a $N \times N$ matrix $\mathbf{A}$, by sampling each element from a uniform distribution: $\mathbf{A}_{ij} \sim \mathcal{U}(0, 1)$
- We performed QR-decomposition on $\mathbf{A}$ to obtain an orthogonal matrix $\mathbf{Q}$, and ensured that the $\mathbf{Q}$ was positive definite by flipping the signs of its elements, if its determinant was negative.
- We fixed a diagonal matrix $\mathbf{D}$ (the values on the diagonal were different depending on the value of $N$).
- We computed the covariance matrix $\Sigma = \mathbf{Q}^T\mathbf{D}\mathbf{Q}$

We generated $L$ different datasets, by sampling different $\mathbf{A}$ and thus constructing different $\Sigma$ for each dataset, and correspondingly sampling $K$ different postsynaptic activity vectors $\mathbf{x} \sim \mathcal{N}(0, \Sigma)$ for each $\Sigma$.

The postsynaptic activity at time $t$ $y_1^t$ was updated using:

$$y_i^t = \sum_{j=1}^{N} \omega_{ij}^t x_j^t \tag{8}$$

where $\omega_{ij}$ is the synaptic weight between the $i$th postsynaptic and $j$th presynaptic neuron. We simulated postsynaptic activity for $T$ timesteps i.e., there were $T$ updates to the postsynaptic activity in one trace, for each of the $K$ presynaptic activity samples, for all $L$ datasets. We kept the presynaptic activity the same for all $T$ timesteps i.e., $x_j^t = x_j^0 = x_j$.

We updated the synaptic weights $\omega_{ij}$ concurrently with the postsynaptic activity, using an update rule $h$, which we denote as $h_\theta$ when the update function is parametrized. Note that the weights were updated using implicit batch learning, i.e. we computed the update $\Delta\omega_{ij} = h(\cdot)$ by averaging over the $K$ different $x_j$ and $y_i^t$ values at teach timestep:

$$\omega_{ij}^{t+1} = \omega_{ij}^t + \frac{\eta}{K}\sum_{k=1}^{K} h(x_j^{(k)}, (y_i^{(k)})^t, \omega_{ij}^t) \tag{9}$$

where $\eta$ is the learning rate. We thus had one synaptic weight trajectory corresponding to batch learning over $K$ different pre- and post-synaptic activity traces for $T$ timesteps. At the start of a simulation over $K$ samples, we set the postsynaptic activity at $t = 0$ to 0, and initialised the synaptic weights randomly ($\omega_{ij}^0 \sim \mathcal{N}(0, 0.1^2)$).

For all experiments described in the main paper, we set $T = 200$, $M = 1$, $K = 100$, and $\eta = 0.1$.

For the proof of principle, and training the local MLP in Fig. 5, and for the test data in Fig. 2-3-4, we set $N = 3$.

For the noisy postsynaptic activity training data in Fig. 3, we set $N = 3$, and modified Eqn 8 as follows:

$$y_i^t = \sum_{j=1}^{N} \omega_{ij}^t x_j^t + \epsilon,\ \epsilon \in \mathbb{R}^M,\ \epsilon \sim \mathcal{N}(0, 0.25^2) \tag{10}$$

For the training data with 39 presynaptic neurons in Fig. 3, we set $N = 39$.

##### B.2 Synaptic update rules

For all training and test data, the groundtruth rule $h$ was Oja's rule (Eqn. 2)

The parametrized update rules $h_\theta$ had the following architecture:

1. **Constrained rule**: $h_\theta(\omega_{ij}^t, x_j, y_i^t) = y_i^t(\theta_1 x_j + \theta_2 y_i^t \omega_{ij}^t)$.
   The parameters of this rule were the scalars $\theta_1$ and $\theta_2$
2. **Local MLP**:
   This was an MLP with 3 fully-connected layers with hidden units $[100, 100, 100]$ and no added bias. Each of the fully-connected layers was followed by a non-linearity: [Sigmoid, Sigmoid]. This MLP was local: it took a 3-dimensional input consisting of one value of the presynaptic activity for one neuron e.g. $j$, the postsynaptic activity of one neuron $i$, and the synaptic weight connecting the two $\omega_{ij}$.
3. **Oja + local MLP**:
   The MLP for this rule had the exact same architecture as for Local MLP. We then added the output of Oja's rule to the output of the MLP.
4. **Semi-global MLP**:
   This MLP had the same architecture as for Local MLP, except that it took a 5D input. The two additional dimensions compared to local MLP were the mean presynaptic activity from $N$ presynaptic neurons, and the mean synaptic weight across $M \times N$ synaptic weights.
5. **Global MLP**:
   This MLP had the same architecture as for the local MLP, but took in a $N + M + N \times M$-D input. This MLP was global: it took the activity of all pre- and postsynaptic neurons in the network as well as all synaptic weights as input, and computed an update for all synaptic weights in a single forward pass. The final layer that reshaped the output of the last fully-connected layer into a $N \times M$-D output.

##### B.3 Discriminator architecture

For the learned rules, the discriminator architecture was as follows:

1. For the constrained rule: 2 convolutional layer with input channels , output channels and kernel size = ($T = 10$, 5, 10) and (5, 3, 3). Each convolutional layer was followed by a 1D MaxPool layer with kernel size 2, and stride 1. These layers were followed by a fully connected layer with 1 hidden unit. Each layer was also followed by a LeakyReLU nonlinearity with slop 0.2. The final fully connected layer was followed by a sigmoid.
2. For all other parametrized rules: 2 convolutional layer with input channels , output channels and kernel size = ($M$, 5, 10) and (5, 1, 10). These layers were followed by 3 fully connected layers with 100 hidden units each. Each layer was also followed by a LeakyReLU nonlinearity with slope 0.2. The final fully connected layer was followed by a sigmoid. For these discriminators, we also used spectral normalisation [36] to stabilize training.

##### B.4 Training details

###### B.4.1 GAN training

To train all GANs, we simulated $L = 500$ datasets, one of which was held out for validation to check if training had converged. The batchsize for all GANs was set to 1 i.e., we passed the GANs one dataset per update of the generator and discriminator networks. We also use gradient norm clipping to stabilise training. From the network with $N = 3, \epsilon = 0$, we additionally generated $L' = 100$ test datasets, on which we performed our post-training analsis. For the networks with 39 presynaptic neurons and with noisy postsynaptic activity, we generated $L' = 20$ test datasets.

For the minimial network, we trained the parameters for 5k epochs, where each epoch consisted of 10 discriminator and 1 generator update. Note that we used only $T = 10$ timesteps of the simulated data to train these networks. We used the Adam optimiser with learning rate 0.0001, $\beta_1 = 0.9$ and $\beta_2 = 0.999$.

For the local MLP, we trained for xx epochs with xx discriminator and xx generator updates in each epoch. We clipped the gradients of both networks to be below xx, and used all $T = 200$ timesteps of the postsynaptic activity.

For the semi-local MLP, we trained for xx epochs with xx discriminator and xx generator updates in each epoch. We clipped the gradients of both networks to be below xx, and used all $T = 200$ timesteps of the postsynaptic activity.

For the global MLP, we trained for xx epochs with xx discriminator and xx generator updates in each epoch. We clipped the gradients of both networks to be below xx, and used all $T = 200$ timesteps of the postsynaptic activity.

For the local MLP on noisy data, we trained for xx epochs with xx discriminator and xx generator updates in each epoch. We clipped the gradients of both networks to be below xx, and used all $T = 200$ timesteps of the postsynaptic activity.

For the local MLP on data with 39 presynaptic neurons, we trained for xx epochs with xx discriminator and xx generator updates in each epoch. We clipped the gradients of both networks to be below xx, and used all $T = 200$ timesteps of the postsynaptic activity.

###### B.4.2 Supervised training

We used the same datasets simulated for the GANs (described in the previous section) to perform supervised training for the local MLP architecture on (a) the noiseless dataset with 3 pre-synaptic neurons (b) the noisy dataset with 3 pre-synaptic neurons and (c) the noiseless dataset with 39 pre-synaptic neurons. For all three datasets, we used the mean-squared error between the generated and groundtruth post-synaptic activity traces, to train the MLPs for xx epochs with gradients clipped to below xx and using all $T = 200$ timesteps of the postsynaptic activity.

###### B.4.3 Training MLP on heatmaps

We also trained the local MLP architecture on the weight update heatmaps (Supp. Fig. 5C). More precisely, we used the pre- and post-synaptic activity values as well as the synaptic weights from the datasets we simulated for the GANs. We randomly sampled these quantities for different timesteps and datasets. We then defined a loss as a mean-squared error between the weight updates from Oja's rule and the local MLP, given the same synaptic weights and pre- and post-synaptic activity values: $L(\theta) = ||\omega_{ij}^{(Oja\ Rule)} - \omega_{ij}^{(Local\ MLP)}||^2 = \sum_{x_j, y_i, \omega_{ij}} ||(x_j y_i - y_i^2 \omega_{ij}) - MLP_\theta(x_j, y_i, \omega_{ij})||^2$. We trained the local MLP for xx epochs, and did not use gradient norm clipping.
